# User-friendly, scalable tools and workflows for single-cell analysis

**DOI:** 10.1101/2020.04.08.032698

**Authors:** P. Moreno, N. Huang, J.R. Manning, S. Mohammed, A. Solovyev, K. Polanski, R. Chazarra, C. Talavera-Lopez, M. Doyle, G. Marnier, B. Grüning, H. Rasche, W. Bacon, Y. Perez-Riverol, M. Haeussler, K.B. Meyer, S. Teichmann, I. Papatheodorou

## Abstract

Single-cell RNA-Seq (scRNA-Seq) data analysis requires expertise in command-line tools, programming languages and scaling on compute infrastructure. As scRNA-Seq becomes widespread, computational pipelines need to be more accessible, simpler and scalable. We introduce an interactive analysis environment for scRNA-Seq, based on Galaxy, with ~70 functions from major single-cell analysis tools, which can be run on compute clusters, cloud providers or single machines, to bring compute to the data in scRNA-Seq.

---

Currently large consortiums such as the Human Cell Atlas[1], the Gut Cell Atlas[2] and the Fly Cell Atlas[3], among others, are generating large amounts of single-cell RNA-seq (scRNA-Seq) data that are or will be publicly accessible. The EMBL-EBI Single Cell Expression Atlas[4] comprises more than 2 million cells from over 14 species and 151 studies; with data readily available to be inspected and re-analyzed. On top of that, many researchers are in the need of analyzing their own scRNA-Seq datasets. This large avalanche of scRNA-Seq data needs to be met by adequate computational infrastructure, analysis tools and workflows to help researchers make the most of this data and re-utilise it.

scRNA-Seq data analysis can be divided into two distinct phases: quantification (producing an expression matrix) and downstream analysis, comprising all subsequent steps including, for example filtering, normalisation, batch correction, marker genes, trajectories analysis and clustering. Downstream analysis functionalities are implemented by a range of different command-line tools and libraries, which in the second case require the user to program in the library’s language of choice. Each of these tools and libraries will excel at certain parts of the analysis and as such best results will be obtained by combining them. Many tools aim to capture a number of different steps in the downstream analysis, but it is unlikely that any given tool will have the best methodology at all stages. This means that a theoretical ‘optimal workflow’ will involve combining elements of different tools, possibly written in different languages (for example R and Python). However the lack of well established exchange formats, the need to switch from one language to the other, setup difficulties and other complications can drive researchers to stick with a single tool for all their analysis steps. This can lead users to miss out on a wider variety of algorithms available for the analysis - particularly where tools change quickly in a field as rapidly developing as single-cell transcriptomics. In the case of clustering tools for instance, Seurat[5] is a highly used framework that requires users to program in R; moving to a competing framework, like Scanpy[6], would require users to also learn Python (as the latter is used as a Python library).

For downstream analysis to be effective, it must be guided by the original experimental design and biological questions, requiring a close involvement of the biological researcher who generated the data. However, using these tools at scale requires computational skills to establish an appropriate computational infrastructure and undertake this analysis, which even for expert bioinformaticians can take significant time and effort. Researchers, frustrated with the complexities of command-line tools or the lack of time to use a tool in a programming library manner, will recourse instead to online tools through web applications that have graphical user interfaces (UIs). This will ease access to a limited range of tools, but will constrain their ability to process large datasets due to upload or bandwidth issues. A better, more scalable solution is the deployment of online applications next to the data (bring the compute to the data).

To alleviate these problems, we present the Hinxton Single Cell Interactive Analysis Portal (HiSCiAp), our solution for integrated analysis of Human Cell Atlas data, although it is generically applicable. HiSCiAp is based on Galaxy, has a number of relevant scRNA-Seq analysis tools, demonstration workflows and data retrieval modules from the EBI Single Cell Atlas and the Human Cell Atlas. It significantly improves access to scRNA-seq data analysis for researchers who are not expert bioinformaticians.

A key feature of HiSCiAp is the ability to integrate tools from different workflows, written in different languages. HiSCiAp breaks monolithic tools into analysis components, providing users with the ability not only to try different competing tool-sets, but also (where possible) integrate them into the same workflows. <u>Supplementary Table 1</u> shows all the tools integrated and the different functional modules in which they were broken; <u>Supplementary Note 1</u> shows the integration of modules from different tools on analysis workflows.

Conventionally, tool interoperability is made possible by exchange file formats being supported by different tools, or through the addition of informatic conversions between them. This adds unnecessary complexity for the researcher, who should be more involved in the decisions of which algorithms to use, rather than having to plumb tools together and negotiate file format issues. In HiSCiAp we support a number of exchange formats, made to be used natively wherever possible by applications, and additional format converters to enable tools that cannot directly read or write them. <u>Supplementary Note 2</u> shows all formats accepted by the different modules, and all conversion modules.

All of the scRNA-Seq tools provided via HiSCiAp are based on command-line interface (CLI) components provided as installable packages via the community-driven Bioconda[7] project. Bioconda produces Docker containers (Biocontainers[8]) automatically for each package. This facilitates software installation and maintenance for researchers and allows reproducibility of analysis across infrastructures and over time (even when ‘current’ software versions have moved on), via strict versioning and dependency management. <u>Supplementary Table 1</u> includes all links to containers and packages.

Direct use of Biocontainers or Bioconda packages on their own for analysis execution still requires some computational expertise, especially if the analysis needs to run on scalable infrastructure. HiSCiAp wraps the Bioconda packages and Biocontainers to provide Galaxy tools and workflows, benefiting from the Galaxy[9] graphical user interface to execute tools, build workflows and share analysis results. Through Galaxy, HiSCiAp can run both on a number of HPC systems, or on cloud providers through the Kubernetes integration[10] with minimal effort, such that the whole setup can be deployed next to the data ingestion points, bringing compute to the data. We have tested HiSCiAp to run on LSF, OpenStack, Amazon Web Services (AWS), Google Cloud Platform and local machines. Furthermore, all of the HiSCiAp Galaxy modules can be installed through the Galaxy ToolShed[11] on any existing Galaxy setup. <u>Supplementary Note 3</u> shows the different installation modalities.

Bioconda and Biocontainers CLI components described above can also be leveraged by other workflow tools with Conda integration (Nextflow and Snakemake, for example) to flexibly generate and execute (also via HPC or cloud) novel workflows without writing new library-specific code, independently of the underlying infrastructure. For direct usage, either through other workflow environments or manually in Linux, these CLI components only require linux command line understanding and no longer the need to use a programming language as a library, reducing entry barriers. These CLI components also facilitate a more uniform use of the underlying analysis software libraries across workflow environments, as otherwise each different workflow environment would need to write its own, often different, library calls to wrap the functionality. For example, besides Galaxy, also Nextflow[12] workflows based on our setup are available in the <u>Supplementary Note 1</u>, and <u>Supplementary Note 2</u> includes all of the generated wrappers.

We have tested this setup with more than 151 scRNA-Seq datasets: HiSCiAp has been used to execute the downstream analysis for all dataset shown on the EMBL-EBI Single Cell Expression Atlas (SCXA) for its past 4 releases, using the Galaxy API to run processes in an automated manner. The workflows used to produce the data for SCXA are publicly available at https://humancellatlas.usegalaxy.eu/. We have also run two training events with a total of 70 assistants using this setup on an <u>HPC system (LSF)</u> and <u>on a private cloud (OpenStack)</u>. HiSCiAp can be deployed as well on Amazon Web Services (AWS) Elastic Kubernetes Service (EKS) and Google Cloud Platform (GCP) Kubernetes, and with minimal additional effort, on any Kubernetes installation.

In summary, HiSCiAp is a suite of components derived from commonly used tools in scRNA-Seq analysis. The setup is built on top of Galaxy, which can be deployed on cloud providers, HPC clusters or existing Galaxy instances, to reduce the software engineering burden on researchers. For consistency with researchers building workflows in other infrastructures, we provide the same components as command-line scripts provided via Bioconda packages and Biocontainers containers. In this way we hope to free a variety of researchers to do single cell downstream analysis at scale.

## Methods

Instructions for running different workflows are available in the <u>Supplementary Note 1</u>. Methods for deploying the setup through the different available modalities are available in the <u>Supplementary Note 4</u>.

## References

1.- Regev A, Teichmann SA, Lander ES, et al. The Human Cell Atlas. Elife. 2017;6:e27041. Published 2017 Dec 5. doi:10.7554/eLife.27041

2.- https://www.gutcellatlas.helmsleytrust.org/

3.- https://flycellatlas.org/

4.- Papatheodorou I, Moreno P, Manning J, et al. Expression Atlas update: from tissues to single cells. Nucleic Acids Res. 2020;48(D1):D77–D83.

5.- Stuart T, Butler A, Hoffman P, et al. Comprehensive Integration of Single-Cell Data. Cell. 2019;177(7):1888–1902.e21.

6.- Wolf FA, Angerer P, Theis FJ. SCANPY: large-scale single-cell gene expression data analysis. Genome Biol. 2018;19(1):15.

7.- Grüning B, Dale R, Sjödin A, et al. Bioconda: sustainable and comprehensive software distribution for the life sciences. Nat Methods. 2018;15(7):475–476.

8.- Da veiga leprevost F, Grüning BA, Alves aflitos S, et al. BioContainers: an open-source and community-driven framework for software standardization. Bioinformatics. 2017;33(16):2580–2582.

9.- Afgan E, Baker D, Batut B, et al. The Galaxy platform for accessible, reproducible and collaborative biomedical analyses: 2018 update. Nucleic Acids Res. 2018;46(W1):W537–W544.

10.- Moreno, P., Pireddu, L., Roger, P., et al. Galaxy-Kubernetes integration: scaling bioinformatics workflows in the cloud. BioRxiv. 2018, 488643.

11.- Blankenberg D, Von kuster G, Bouvier E, et al. Dissemination of scientific software with Galaxy ToolShed. Genome Biol. 2014;15(2):403.

12.- Di tommaso P, Chatzou M, Floden EW, Barja PP, Palumbo E, Notredame C. Nextflow enables reproducible computational workflows. Nat Biotechnol. 2017;35(4):316–319.

