## Supplementary material for "User-friendly, scalable tools and workflows for single-cell analysis": Supplental Note 1

### Available workflows

Tools can be run in any workflow environment, but here we focus on Galaxy and Nextflow examples.

The most up-to-date version of this supplementary material is available here.

#### Galaxy

Table S3 below gives a summary of example workflows where different tools are deployed in Galaxy. Note in particular the use intermediate exchange formats or conversions to connect different tools. The sections below explain in more detail each of the workflows.

To run the Galaxy workflows, the easiest is to create an account in https://humancellatlas.usegalaxy.eu/ (if you already have an account in https://usegalaxy.eu/, it is the same account). To create that account go to https://usegalaxy.eu/login and click on `Register here`.

Table S3: Workflows with links to Galaxy instance and to their source definition, assigned to one or more of the relevant analysis areas: Clustering (C), Differential expression/Marker detection (DE-MD), Trajectories (T), Cell type alignment (CT) and Dimensionality reduction (DR). All the workflow source files are available here.

| Workflow | Description | Analysis areas |
|---|---|---|
| Atlas-Scanpy-CellBrowser | Retrieves data from Single Cell Expression Atlas for a given accession, filters, normalise, clusterise, marker genes and calculate dimensional reduction with Scanpy. Visualise interactively with UCSC CellBrowser | C DR DE IV |
| Atlas-Seurat-CellBrowser | Retrieves data from Single Cell Expression Atlas for a given accession, filters, normalise, clusterise, marker genes and calculate dimensional reduction with Seurat. Visualise interactively with UCSC CellBrowser | C DR DE IV |
| SC-Atlas-Production | Filtering, normalization, clustering, marker genes and dimensionality reduction used to process every dataset shown in the Single Cell Expression Atlas release 5 and 6 | C DR DE |

| | | |
|---|---|---|
| HCA-Scanpy-CellBrowser | Retrieves data from Human Cell Atlas for a given accession, filters, normalise, clusterise, marker genes and calculate dimensional reduction with Scanpy. Visualise interactively with UCSC CellBrowser | C DR DE IV |
| Atlas-Scanpy-SCCAF* | Retrieves data from Single Cell Expression Atlas, preprocess and clusterise with Scanpy at two different resolutions and search for best clustering with SCCAF after batch correcting. | C DR DE BC |
| Atlas-Scanpy-SCMap* | Retrieves data from Single Cell Expression Atlas, preprocess and map to an SCMap cell index | C CT |

#### Atlas Scanpy CellBrowser

This workflow (available here at the Human Cell Atlas Galaxy instance) retrieves quantified data matrix
in 10x format from the Single Cell Expression Atlas through an accession number. The data is transformed into AnnData, the format used by Scanpy, and then a downstream analysis is carried out with Scanpy. This analysis first filters the matrix by cells and genes attributes, normalises it, find variable genes, scale it, run PCA, compute the k-nn graph, find clusters, marker genes and calculate tSNE and UMAP projections. The final result is passed to UCSC CellBrowser for interactive visualisation based on the calculated embeddings. Each of the steps provides a number of parameters to adapt for particular uses and datasets.

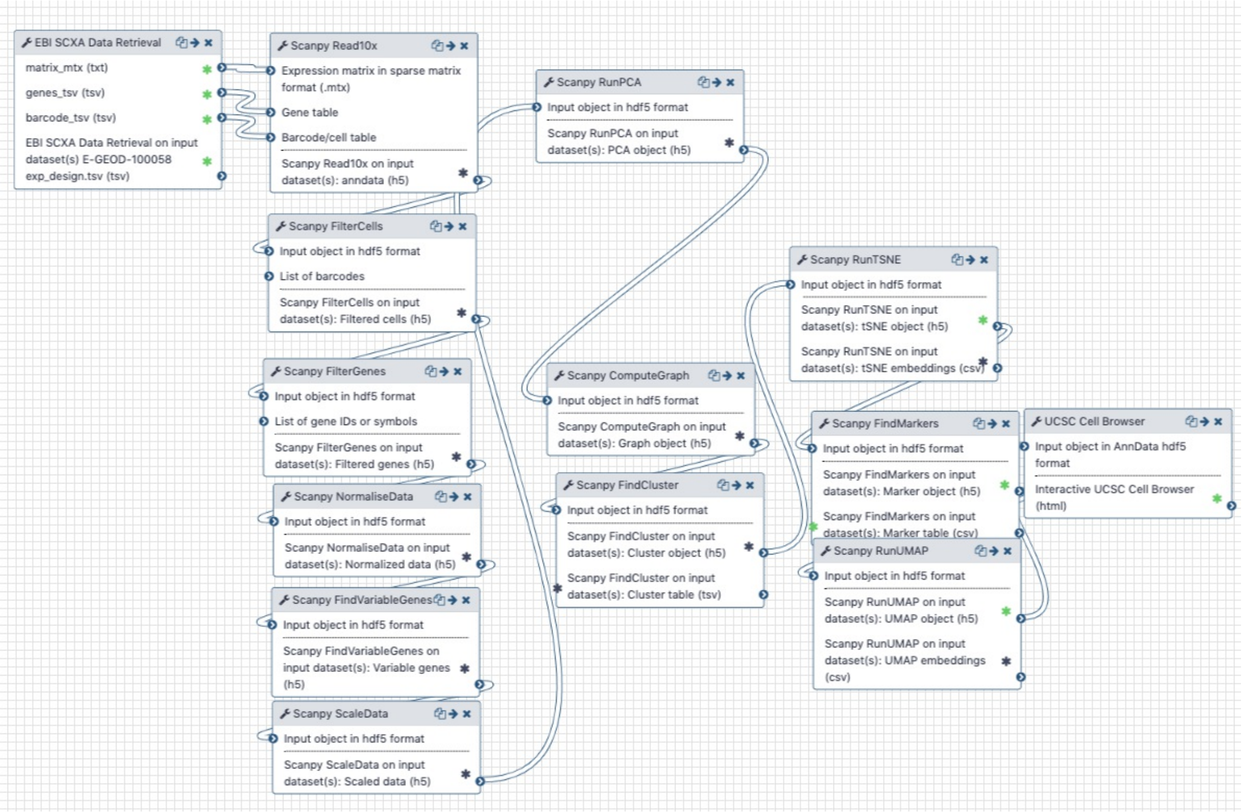


To run this workflow:

- Import the workflow: Click here and then click on the + (plus) button next to "About this workflow", to import it.
- On the next window click on `Start using this workflow`.
- From the Workflows view, locate the new workflow (it will be called "Imported: EBI-Single-Cell-ExpAtlas-Scanpy-CellBrowser") and click on `Run` or `Edit`. `Edit` will allow you to make changes on the workflow.
- If pressed `Run`, on the next screen click on the pencil box next to `SC-Atlas experiment accession` to set the accession of the EMBL-EBI Single Cell Expression dataset.
- Click on the blue button `Run Workflow`. Results will start to appear on the right panel (History).

### Atlas Seurat CellBrowser

This workflow, is analogous to the previous one, but uses instead of Scanpy the downstream analysis steps of Seurat.

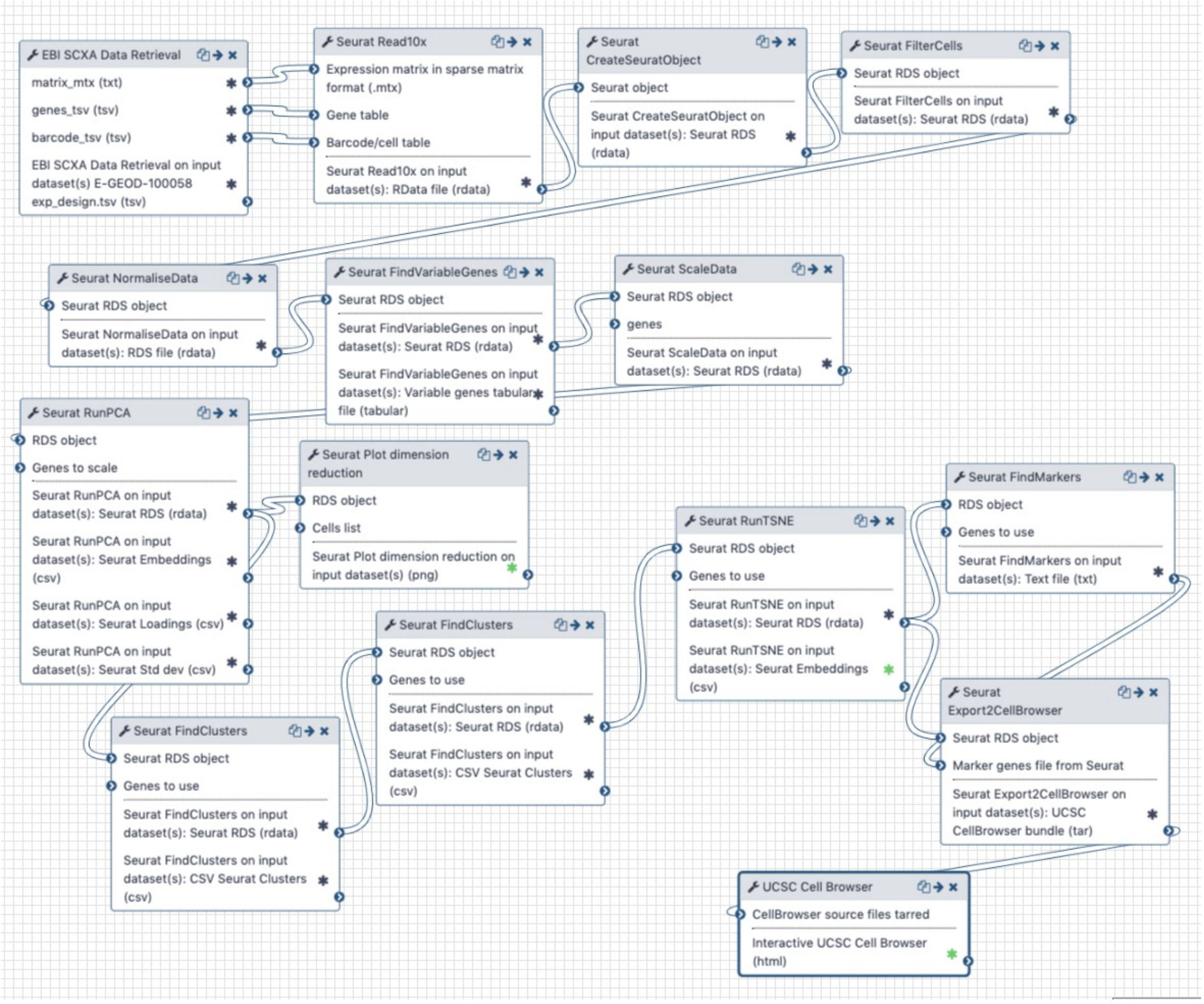


To run this workflow:

- Import the workflow: Click here and then click on the + (plus) button next to "About this workflow", to import it.
- On the next window click on `Start using this workflow`.
- From the Workflows view, locate the new workflow (it will be called "Imported: Atlas-Seurat-CellBrowser") and click on `Run` or `Edit`. `Edit` will allow you to make changes on the workflow.
- If pressed `Run`, on the next screen click on the pencil box next to `SC-Atlas experiment accession` to set the accession of the EMBL-EBI Single Cell Expression dataset.
- Click on the blue button `Run Workflow`. Results will start to appear on the right panel (History).

### SC Atlas Production

This workflow was used for releases 4, 5 and 6 of the EBI Single Cell Expression Atlas. Currently it relies mostly on Scanpy steps, but as interoperability improves, some of the analysis steps could be changed to other software providers. It distributes generation of tSNE plots across different persplexity values and clusters through a set of resolutions. This workflow mostly generates text files that are loaded into databases and indexes for expression atlas, so no interactive modules or plots are generated, and is provided to enable reproducibility of our results.

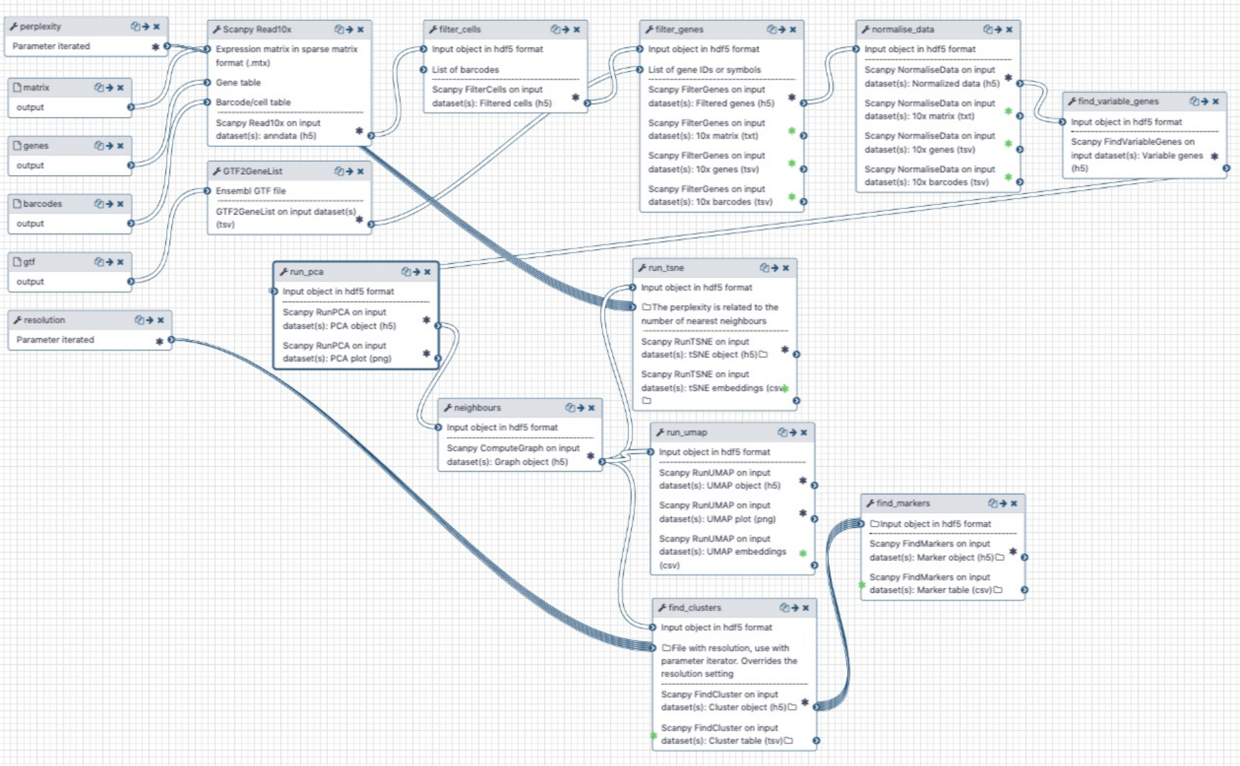


To run this workflow:

- Import the workflow: Click here
- Import an example history with data: Click here and then click on the + (plus) button next to "About this History".
- From the Workflows view, locate the new workflow (it will be called "Imported: EBI Single Cell Expression Atlas Scanpy Prod 1.3") and click on `Run` or `Edit`. `Edit` will allow you to make changes on the workflow.
- If pressed `Run`, make sure that the matrix input points to the Matrix file in the history, the genes input to the `E-HCAD-5.genes` file in the history, the barcode input to the `E-HCAD-5.barcodes` file in the history and the GTF input to the GTF file available in the history.
- Click on the blue button `Run Workflow`. Results will start to appear on the right panel (History).

Datasets can be downloaded through the EBI SCXA Data Retrieval or from the EBI FTP. For the species of the studies, retrieve the GTF (no chr) file from ENSEMBL. In the future, the GTF download will be facilitated.

### Human cell atlas Scanpy CellBrowser

Equivalent to the first workflow using EBI Single Cell Atlas downloader, Scanpy modules and CellBrowser, but using the Human Cell Atlas matrix service as input.

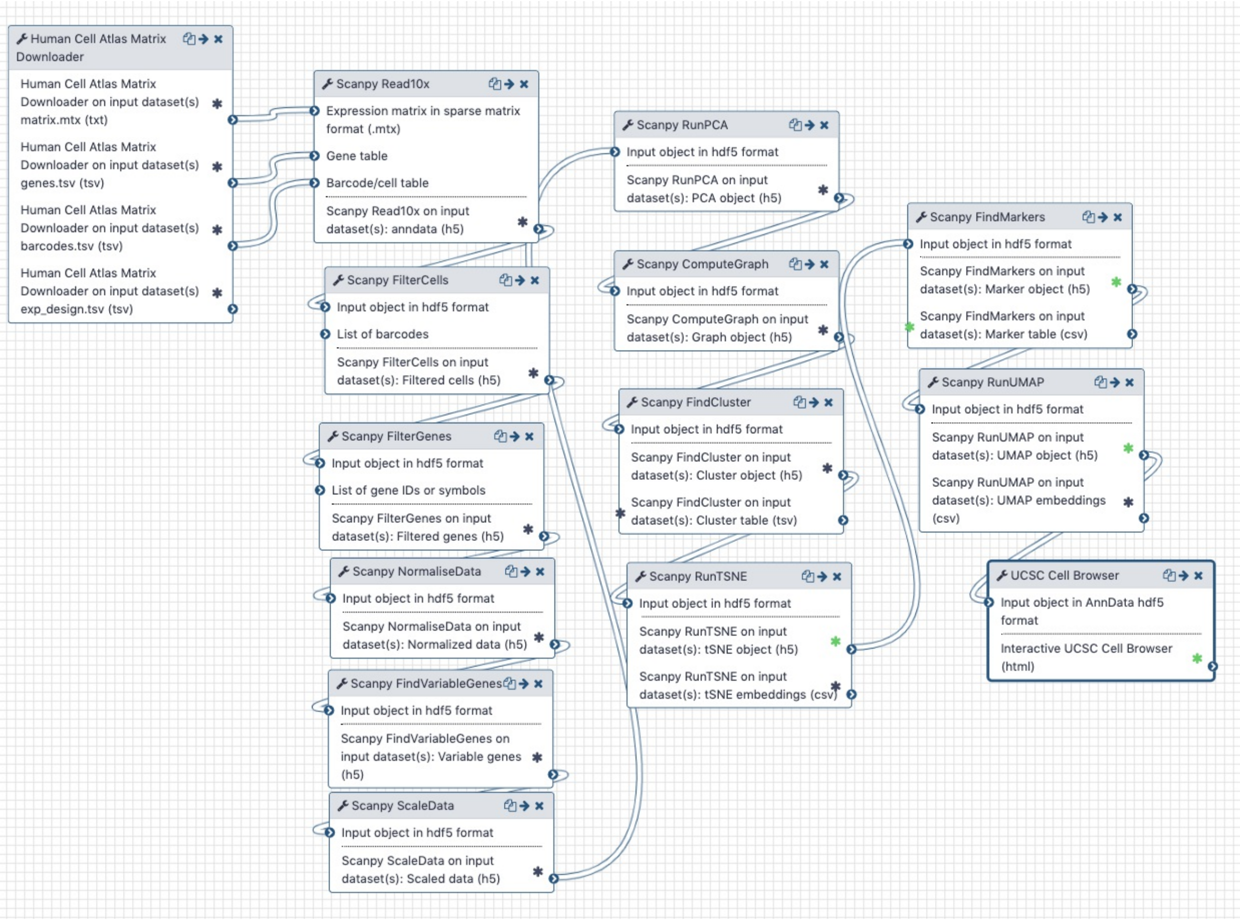


To run this workflow:

- Import the workflow: Click here and then click on the + (plus) button next to "About this workflow", to import it.
- On the next window click on `Start using this workflow`.
- From the Workflows view, locate the new workflow (it will be called "imported: HumanCellAtlas-Scanpy-CellBrowser") and click on `Run` or `Edit`. `Edit` will allow you to make changes on the workflow.
- If pressed `Run`, on the next screen click on the pencil box next to `Human Cell Atlas project name/label/UUID` and set the name, label, or ID for the desired project, which can be found at the HCA data browser, and select the desired matrix format (Matrix Market or Loom). You can leave the default example if you want.
- Click on the blue button `Run Workflow`. Results will start to appear on the right panel (History), unless that you selected to create a new history with the results.

##### Atlas Scanpy SCCAF

This workflow helps in the decision of the number of clusters to pick for a dataset through the use of the SCCAF tool. It will run Scanpy with two different resolution values, a high resolution and a low resolution, and through a machine learning process provide an assessment of adequate clustering in between these settings to prefer for the dataset at hand.

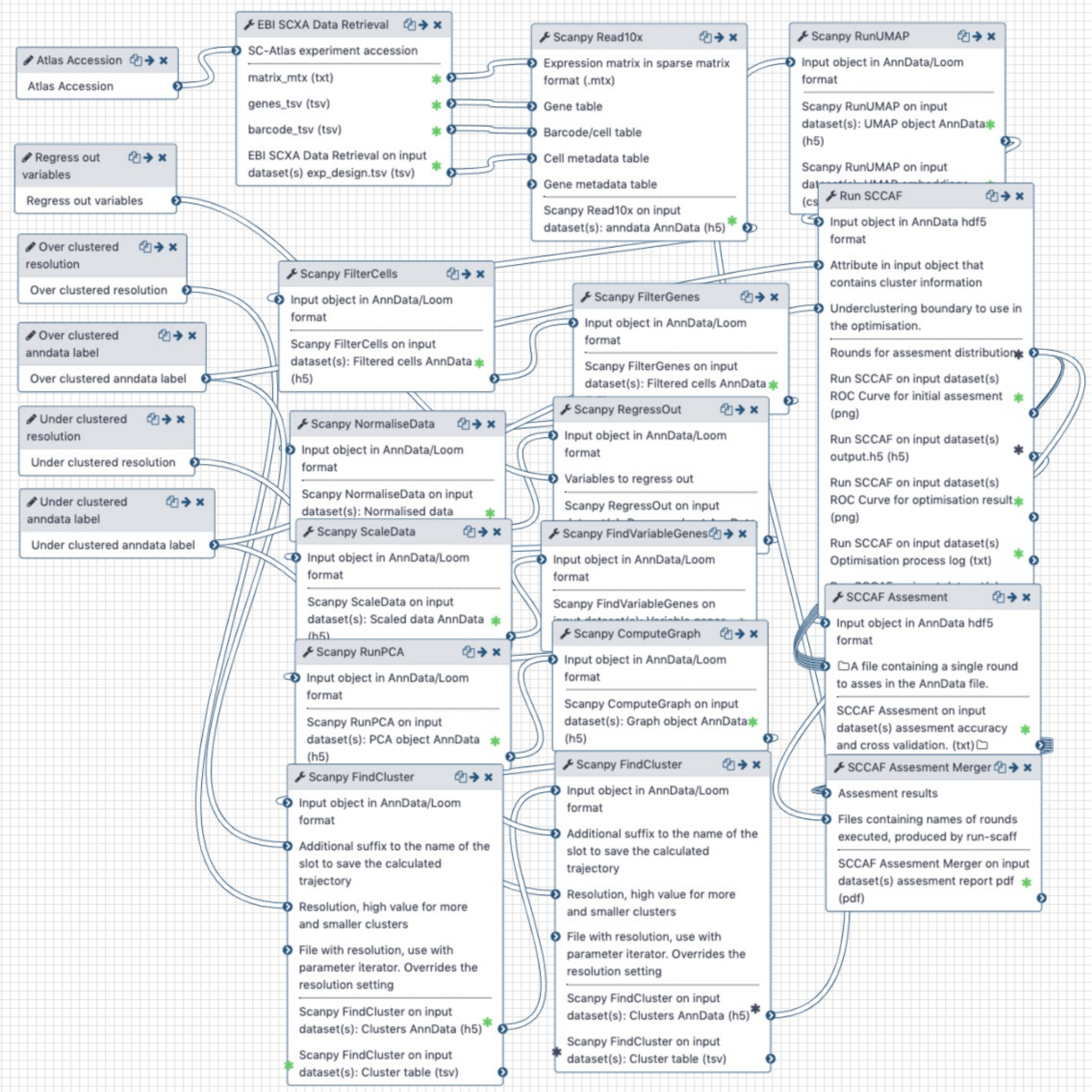


To run this workflow:

- Import the workflow: Click here and then click on the + (plus) button next to "About this workflow", to import it.
- On the next window click on `Start using this workflow`.
- From the Workflows view, locate the new workflow (it will be called "imported: EBI Single Cell Expression Atlas Scanpy SCCAF") and click on `Run` or `Edit`. `Edit` will allow you to make changes on the workflow.
- If pressed `Run`, on the next screen click on the pencil box next to `SC-Atlas experiment`

`accession` to set the accession of the EMBL-EBI Single Cell Expression dataset. You can use for instance `E-MTAB-7195`. Set as well the over and under clustered resolutions (to fractions below 1, under clustered smaller and over clustered bigger) and the over and under clustered labels (just two labels with no spaces to name them).
- Click on the blue button `Run Workflow`. Results will start to appear on the right panel (History), unless that you selected to create a new history with the results.

#### Atlas Scanpy SCMap

This workflow does the downstream analysis with Scanpy for one Atlas dataset and projects it through SCMap at the level of cells and clusters to cell and clusters indexes provided in a format that SCMap can read. The scanpy analysis is exported as Loom file and transformed into SingleCellExperiment by the SCEasy conversion module.

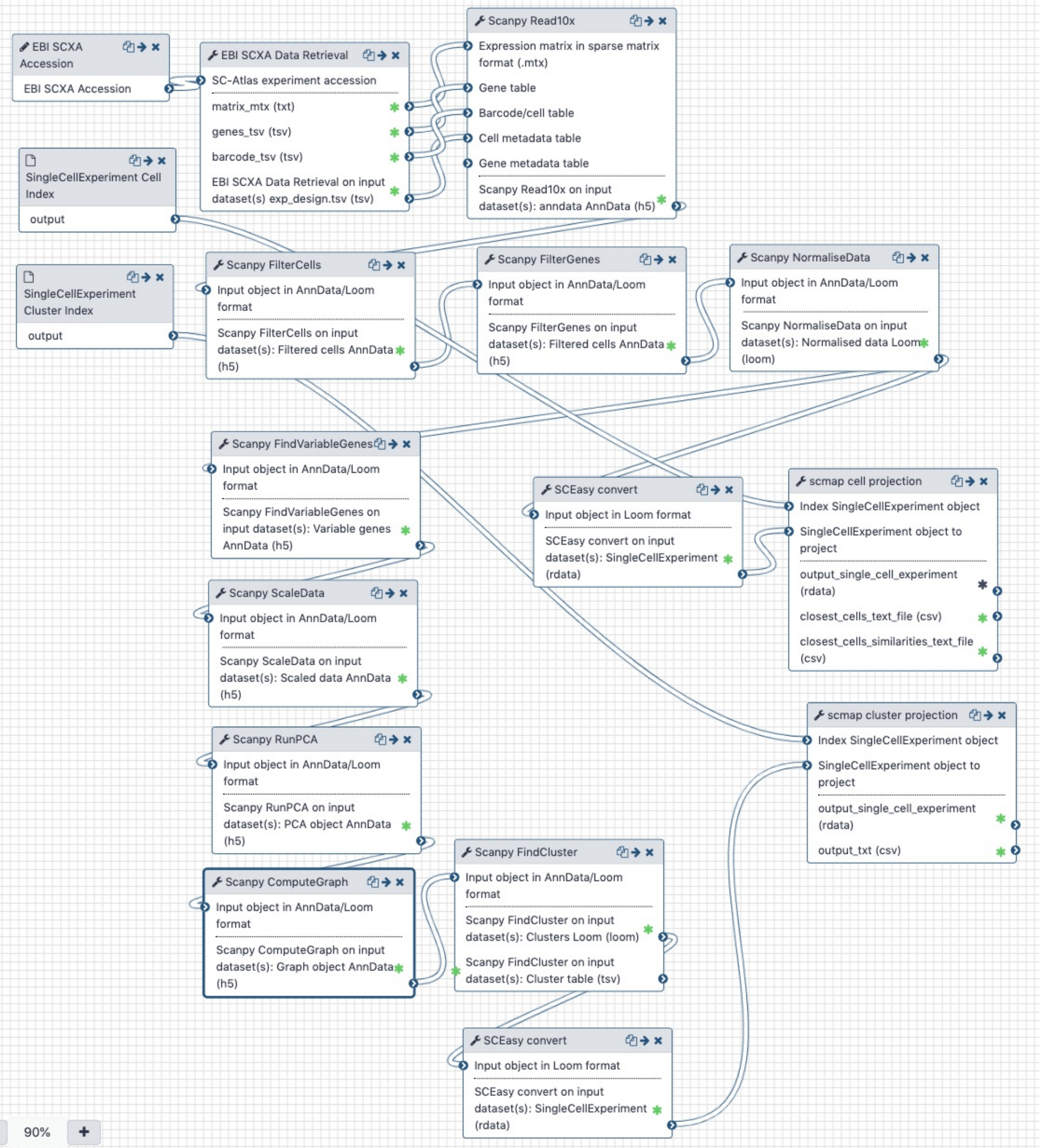


### Nextflow

Table S4 below shows examples where tools are deployed in Nextflow workflows. Areas: Cell type prediction (CTP), QC/ preprocessing (QC/P).

| Workflow | Description | Analysis areas |
|---|---|---|
| scmap-prod-workflow | Predict cell types using scmap, incorporating tools of | CTP |

| | | |
|---|---|---|
| | the DropletUtils CLI | |
| scpred-prod-workflow | Predict cell types using scPred, incorporating tools of the DropletUtils CLI | CTP |
| garnett-prod-workflow | Predict cell types using Garnett, incorporating tools of the Monocle CLI | CTP |
| droplet-quantification-workflow | Used by the EBI Gene Expression team for quantifying droplet single-cell experiments, incorporating tools of the DropletUtils CLI | QC/P |
