## Supplemental Note 2 for "User-friendly, scalable tools and workflows for single-cell analysis"

### Available Tools

HSCiAP relies on published analysis tools for single downstream analysis, starting from a quantified matrix (usually in 10x format). In the process of wrapping those tools, we have made an effort to isolate its main analysis steps, towards an scenario where different analysis components from each tool can be used interchangeably provided by that their adherence to exchange formats, or intermediate converters, allow so. Tables contain all the tools currently wrapped, with links to their sorce codes and our wrappers.
The following section goes through each tool and the steps that we extracted.

The most up-to-date version of this supplementary material will be available here.

### Scanpy

From its GitHub repository, Scanpy is defined as:

> Scanpy is a scalable toolkit for analyzing single-cell gene expression data built jointly with anndata. It includes preprocessing, visualization, clustering, trajectory inference and differential expression testing. The Python-based implementation efficiently deals with datasets of more than one million cells.

Scanpy is used as a Python library, so Python code needs to be written to use it directly. We have written
a CLI component named scanpy-scripts which allows its usage from the command line, avoiding the need
of having to write Python code to use it for the wrapped functionality. The section Scanpy Steps available
below details all the steps wrapped.

#### Scanpy Exchange formats

- 10x: initial reading and as output in some steps.
- AnnData: read/write through most modules.
- Loom: read/write through most modules.

#### Scanpy modules available

Table 1 below details all modules available as both a CLI component and as a Galaxy wrapper. The name of the module links to an active Galaxy instance where that can be used, and TS provides a second link to the module in the Galaxy Toolshed.
Each module is linked to one of the cli-layers to one or more of the relevant analysis areas: Clustering (C), Differential expression/Marker detection (DE-MD), Trajectories (T), Cell type alignment (CT), quality control (QC) and Dimensionality reduction (DR).

| Module | Description | cli-layer | Analysis areas |
|---|---|---|---|
| scanpy_filter_cells TS | based on counts and numbers of genes expressed | scanpy-scripts | QC |
| scanpy_filter_genes TS | based on counts and numbers of cells expressed | scanpy-scripts | QC |
| scanpy_find_cluster TS | based on community detection on KNN graph | scanpy-scripts | C |
| scanpy_find_markers TS | to find differentially expressed genes between groups | scanpy-scripts | DE-MD |
| scanpy_find_variable_genes TS | based on normalised dispersion of expression | scanpy-scripts | P |
| scanpy_compute_graph TS | to derive kNN graph | scanpy-scripts | C T |
| scanpy_normalise_data TS | to make all cells having the same total expression | scanpy-scripts | P |
| scanpy_parameter_iterator TS | produce an iteration over a defined parameter | scanpy-scripts | |
| scanpy_plot_embed TS | visualise cell embeddings | scanpy-scripts | C |
| scanpy_plot_trajectory TS | visualise cell trajectories | scanpy-scripts | T |

| | | | |
|---|---|---|---|
| scanpy_read_10x[TS] | into hdf5 object handled by scanpy | scanpy-scripts | |
| scanpy_regress_variable[TS] | variables that might introduce batch effect | scanpy-scripts | P |
| scanpy_run_diffmap[TS] | calculate diffusion components | scanpy-scripts | T |
| scanpy_run_dpt[TS] | diffusion pseudotime inference | scanpy-scripts | T |
| scanpy_run_fdg[TS] | visualise cell clusters using force-directed graph | scanpy-scripts | |
| scanpy_run_paga[TS] | trajectory inference | scanpy-scripts | T |
| scanpy_run_pca[TS] | for dimensionality reduction | scanpy-scripts | DR |
| scanpy_run_tsne[TS] | visualise cell clusters using tSNE | scanpy-scripts | DR |
| scanpy_run_umap[TS] | visualise cell clusters using UMAP | scanpy-scripts | DR |
| scanpy_scale_data[TS] | to make expression variance the same for all genes | scanpy-scripts | P |

#### Examples of Workflows using Scanpy

### Seurat

According to its website, Seurat is:

> …an R package designed for QC, analysis, and exploration of single-cell RNA-seq data. Seurat aims to enable users to identify and interpret sources of heterogeneity from single-cell transcriptomic measurements, and to integrate diverse types of single-

> cell data.

Seurat, as any R package, requires the user to write R code in order to use it to analyze data. We have written a CLI component, Seurat scripts, which exposes relevant functionality through a command line interface, enabling its use without having to write R code.

#### Seurat exchange formats

Seurat version 3 wrapped supports the following exchange formats:

- 10x (input)
- Loom (input and output in most steps)
- AnnData (input for most steps)
- SingleCellExperiment (input and output for most steps) as an RDS R object.
- Seurat native format as an RDS R object.

#### Seurat modules available

Table 2 below details all modules available as both a CLI component and as a Galaxy wrapper. The name of the module links to an active Galaxy instance where that can be used, and TS provides a second link to the module in the Galaxy Toolshed.
Each module is linked to one of the cli-layers to one or more of the relevant analysis areas: Clustering (C), Differential expression/Marker detection (DE-MD), Trajectories (T), Cell type alignment (CT), quality control (QC) and Dimensionality reduction (DR).

| Module | Description | cli-layer | Analysis areas |
|---|---|---|---|
| seurat_dim_plot TS | graphs the output of a dimensional reduction technique (PCA by default). Cells are colored by their identity class. | seurat-scripts | DR |
| seurat_export_cellbrowser TS | produces files for UCSC CellBrowser import. | seurat-scripts | |
| seurat_filter_cells TS | filter cells in a Seurat object | seurat-scripts | |
| | | seurat- | |

| | | | |
|---|---|---|---|
| seurat_find_clusters[TS] | find clusters of cells | scripts | C |
| seurat_find_markers[TS] | find markers (differentially expressed genes) | seurat-scripts | DE-MD |
| seurat_find_variable_genes[TS] | identify variable genes | seurat-scripts | C DE-MD |
| seurat_normalise_data[TS] | normalise data | seurat-scripts | C |
| seurat_read10x[TS] | Loads 10x data into a serialized seurat R object | seurat-scripts | |
| seurat_run_pca[TS] | run a PCA dimensionality reduction | seurat-scripts | DR |
| seurat_run_tsne[TS] | run t-SNE dimensionality reduction | seurat-scripts | C DR |
| seurat_scale_data[TS] | scale and center genes | seurat-scripts | C |
| seurat_find_neighbours[TS] | constructs a Shared Nearest Neighbor (SNN) Graph | seurat-scripts | C |

# SC3

According to its website, Single-Cell Consensus Clustering (SC3) is:

> …a tool for unsupervised clustering of scRNA-seq data. SC3 achieves high accuracy and robustness by consistently integrating different clustering solutions through a consensus approach. An interactive graphical implementation makes SC3 accessible to a wide audience of users. In addition, SC3 also aids biological interpretation by identifying marker genes, differentially expressed genes and outlier cells. A manuscript describing SC3 in details is published in Nature Methods.

SC3, as any R package, requires the user to write R code in order to use it to analyse data. We have written a CLI component, SC3 scripts, which exposes relevant functionality through a command line interface, enabling its use without having to write R code.

#### SC3 exchange formats

SC3 wrapped supports the following exchange formats:

- 10x (input)
- SingleCellExperiment (input and output for most steps) as an RDS R object.

#### SC3 modules available

Table 3 below details all modules available as both a CLI component and as a Galaxy wrapper. The name of the module links to an active Galaxy instance where that can be used, and TS provides a second link to the module in the Galaxy Toolshed.
Each module is linked to one of the cli-layers to one or more of the relevant analysis areas: Clustering (C), Differential expression/Marker detection (DE-MD), Trajectories (T), Cell type alignment (CT), quality control (QC) and Dimensionality reduction (DR).

| Module | Description | cli-layer | Analysis areas |
|---|---|---|---|
| sc3_calc_biology TS | calculates DE genes, marker genes and cell outliers | sc3-scripts | DE-MD |
| sc3_calc_consens TS | from multiple runs of k-means clustering | sc3-scripts | C |
| sc3_calc_dists TS | between cells | sc3-scripts | |
| sc3_calc_transfs TS | of distances using PCA and graph Laplacian | sc3-scripts | C |
| sc3_estimate_k TS | the number of clusters for k-means clustering | sc3-scripts | C |
| sc3_kmeans TS | perform k-means clustering | sc3-scripts | C |
| sc3_prepare TS | a sc3 SingleCellExperiment object | sc3-scripts | |

### Scater

According to its website, Scater:

> …contains tools to help with the analysis of single-cell transcriptomic data, focusing on low-level steps such as quality control, normalization and visualization. It is based on the SingleCellExperiment class (from the SingleCellExperiment package), and thus is interoperable with many other Bioconductor packages such as scran, batchelor and iSEE.

Scater, as any R package, requires the user to write R code in order to use it to analyse data. We have written a CLI component, scater scripts, which exposes relevant functionality through a command line interface, enabling its use without having to write R code.

### Scater exchange formats

Scater wrapped supports the following exchange formats:

- 10x (input)
- SingleCellExperiment (input and output for most steps) as an RDS R object.

### Scater modules available

Table 4 below details all modules available as both a CLI component and as a Galaxy wrapper. The name of the module links to an active Galaxy instance where that can be used, and TS provides a second link to the module in the Galaxy Toolshed.
Each module is linked to one of the cli-layers to one or more of the relevant analysis areas: Clustering (C), Differential expression/Marker detection (DE-MD), Trajectories (T), Cell type alignment (CT), quality control (QC) and Dimensionality reduction (DR).

| Module | Description | cli-layer | Analysis areas |
|---|---|---|---|
| scater_calculate_cpm[TS] | from raw counts | scater-scripts | |
| scater_calculate_qc_metrics[TS] | based on expression values and experiment information | scater-scripts | |
| scater_filter[TS] | cells and genes based on pre-calculated stats and | scater- | |

| | | | |
|---|---|---|---|
| | QC metrics | scripts | |
| scater_is_outlier TS | cells based on expression metrics | scater-scripts | |
| scater_normalize TS | expression values by library size in log2 scale | scater-scripts | C |
| scater_read_10x_results TS | Loads 10x data into a serialized scater R object | scater-scripts | |

### Monocle3

According to its website, Monocle3:

> …can help you perform three main types of analysis:
>
> - Clustering, classifying, and counting cells. Single-cell RNA-Seq experiments allow you to discover new (and possibly rare) subtypes of cells. Monocle 3 helps you identify them.
> - Constructing single-cell trajectories. In development, disease, and throughout life, cells transition from one state to another. Monocle 3 helps you discover these transitions.
> - Differential expression analysis. Characterizing new cell types and states begins with comparisons to other, better understood cells. Monocle 3 includes a sophisticated, but easy-to-use system for differential expression.

Monocle3, as any R package, requires the user to write R code in order to use it to analyse data. We have written a CLI component, monocle3 scripts, which exposes relevant functionality through a command line interface, enabling its use without having to write R code.

#### Monocle3 exchange formats

Monocle3 wrapped supports the following exchange formats:

- TSV, CSV or RDS Gene expression matrix, plus metadata

#### Monocle3 modules available

Table 5 below details all modules available as both a CLI component and as a Galaxy wrapper. The name of the module links to an active Galaxy instance where that can be used, and TS provides a second link to the module in the Galaxy Toolshed.
Each module is linked to one of the cli-layers to one or more of the relevant analysis areas: Clustering (C), Differential expression/Marker detection (DE-MD), Trajectories (T), Cell type alignment (CT), quality control (QC) and Dimensionality reduction (DR).

| Module | Description | cli-layer | Analysis areas |
|---|---|---|---|
| monocle3_create[TS] | a Monocle3 object from input data | monocle-scripts | |
| monocle3_diffExp[TS] | of genes along a trajectory | monocle-scripts | DE-MD |
| monocle3_learnGraph[TS] | between cells in dimensionality reduced space | monocle-scripts | T |
| monocle3_orderCells[TS] | along trajectories | monocle-scripts | T |
| monocle3_partition[TS] | of cells into groups | monocle-scripts | C T |
| monocle3_plotCells[TS] | in the reduced dimensionality space | monocle-scripts | T |
| monocle3_preprocess[TS] | a Monocle3 object to an initially dimensionally reduced space | monocle-scripts | P |
| monocle3_reduceDim[TS] | for downstream analysis | monocle-scripts | DR |

### SCMap

According to its website, SCMap is:

> …a method for projecting cells from a scRNA-seq experiment onto the cell-types or individual cells identified in other experiments (the application can be run for free, without restrictions, from http://www.hemberg-lab.cloud/scmap).

SCMap, as any R package, requires the user to write R code in order to use it to analyse data. We have written a CLI component, scmap scripts, which exposes relevant functionality through a command line interface, enabling its use without having to write R code.

### SCMap exchange formats

SCMap wrapped supports the following exchange formats:

- SingleCellExperiment (input/output)
- Tab-separated text file (output)

### SCMap modules available

Table 6 below details all modules available as both a CLI component and as a Galaxy wrapper. The name of the module links to an active Galaxy instance where that can be used, and TS provides a second link to the module in the Galaxy Toolshed.
Each module is linked to one of the cli-layers to one or more of the relevant analysis areas: Clustering (C), Differential expression/Marker detection (DE-MD), Trajectories (T), Cell type alignment (CT), quality control (QC) and Dimensionality reduction (DR).

| Module | Description | cli-layer | Analysis areas |
|---|---|---|---|
| scmap_get_std_output TS | Create final output in standard format to allow for downstream analysis of predicted labels by tools of the EBI gene expression group's cell-types-analysis package | scmap-cli | |
| scmap_index_cell TS | creates a cell index for a dataset to enable fast approximate nearest neighbour search | scmap-cli | CT |
| scmap_index_cluster TS | calculates centroids of each cell type and merges them into a single table | scmap-cli | CT |
| | searches each cell in a query dataset for the nearest | scmap- | |

| | | | |
|---|---|---|---|
| scmap_scmap_cell[TS] | neighbours by cosine distance within a collection of reference datasets. | cli | CT |
| scmap_scmap_cluster[TS] | projects one dataset to another | scmap-cli | CT |
| scmap_select_features[TS] | finds the most informative features (genes/transcripts) for projection. | scmap-cli | CT |

### SCCAF

According to its website, Single Cell Clustering Assessment Framework (SCCAF) is:

> …a novel method for automated identification of putative cell types from single cell RNA-seq (scRNA-seq) data. By iteratively applying clustering and a machine learning approach to gene expression profiles of a given set of cells, SCCAF simultaneously identifies distinct cell groups and a weighted list of feature genes for each group. The feature genes, which are overexpressed in the particular cell group, jointly discriminate the given cell group from other cells. Each such group of cells corresponds to a putative cell type or state, characterised by the feature genes as markers.

SCCAF provides a CLI component, which exposes relevant functionality through a command line interface, enabling its use without having to write Python code.

#### SCCAF exchange formats

SCCAF wrapped supports the following exchange formats:

- AnnData (input and output)
- Loom (input)

#### SCCAF modules available

Table 7 below details all modules available as both a CLI component and as a Galaxy wrapper. The name of the module links to an active Galaxy instance where that can be used, and TS provides a second link to the module in the Galaxy Toolshed.
Each module is linked to one of the cli-layers to one or more of the relevant analysis areas:

Clustering (C), Differential expression/Marker detection (DE-MD), Trajectories (T), Cell type alignment (CT), quality control (QC) and Dimensionality reduction (DR).

| Module | Description | cli-layer | Analysis areas |
|---|---|---|---|
| run_sccaf[TS] | to assess and optimise clustering | None | C |
| sccaf_asses[TS] | runs an assesment of an SCCAF optimisation result or an existing clustering. | None | |
| sccaf_asses_merger[TS] | brings together distributed assesments. | None | C |
| sccaf_regress_out[TS] | with multiple categorical keys on an AnnData object. | None | |

### Data retrieval

We provide two modules for data retrieval of expression matrices, one from EMBL-EBI Single Cell Expression Atlas and one from the Human Cell Atlas (HCA) Matrix service. They both require an study identifier (or description as well in the case of the HCA matrix service), and produce output in 10x format.

Table 8 below details all modules available as both a CLI component and as a Galaxy wrapper. The name of the module links to an active Galaxy instance where that can be used, and TS provides a second link to the module in the Galaxy Toolshed.
Each module is linked to one of the cli-layers to one or more of the relevant analysis areas: Clustering (C), Differential expression/Marker detection (DE-MD), Trajectories (T), Cell type alignment (CT), quality control (QC) and Dimensionality reduction (DR).

| Module | Description | cli-layer | Analysis areas |
|---|---|---|---|
| hca_matrix_downloader[TS] | retrieves expression matrices and metadata from the Human Cell Atlas. | hca-matrix-downloader | |
| | Retrieves expression | | |

| | | |
|---|---|---|
| retrieve_scxa[TS] | matrixes and metadata from EBI Single Cell Expression Atlas (SCXA) | None |

### Format conversion

SCEASY provides the following formats conversion:

| From | To | Comments |
|---|---|---|
| Seurat | AnnData | |
| Seurat | SingleCellExperiment | |
| SingleCellExperiment | AnnData | |
| Loom | AnnData | |
| Loom | SingleCellExperiment | |

Table 9 below details all modules available as both a CLI component and as a Galaxy wrapper. The name of the module links to an active Galaxy instance where that can be used, and TS provides a second link to the module in the Galaxy Toolshed.
Each module is linked to one of the cli-layers to one or more of the relevant analysis areas: Clustering (C), Differential expression/Marker detection (DE-MD), Trajectories (T), Cell type alignment (CT), quality control (QC) and Dimensionality reduction (DR).

| Module | Description | cli-layer | Analysis areas |
|---|---|---|---|
| sceasy_convert[TS] | a data object between formats | None | C |

### Visualisation

According to its website, UCSC CellBrowser is:

> a viewer for single cell data. You can click on and hover over cells to get meta information, search for genes to color on and click clusters to show cluster-specific marker genes.
> To look at a list of selected single cell datasets, see http://cells.ucsc.edu

UCSC CellBrowser provides a CLI component, which exposes relevant functionality through a command line interface, enabling its use without having to write Python code.

#### Exchange formats

UCSC CellBrowser supports the following exchange formats:

- AnnData (input)
- Seurat archive (input)
- Loom (input)

#### Modules available

Table 10 below details all modules available as both a CLI component and as a Galaxy wrapper. The name of the module links to an active Galaxy instance where that can be used, and TS provides a second link to the module in the Galaxy Toolshed.
Each module is linked to one of the cli-layers to one or more of the relevant analysis areas: Clustering (C), Differential expression/Marker detection (DE-MD), Trajectories (T), Cell type alignment (CT), quality control (QC) and Dimensionality reduction (DR).

| Module | Description | cli-layer | Analysis areas |
|---|---|---|---|
| ucsc_cell_browser TS | displays single-cell clusterized data in an interactive web application. | None | |

### Garnett

From its website, Garnett allows users to:

> - Train cell type classifiers: Use your single-cell RNA-seq data to build a cell type classifier.
> - Classify your cells: Use pre-trained classifiers to identify cell types in your data

Garnett is used as an R library, so R code needs to be written to use it directly. We have written
a CLI component named garnett-cli which allows its usage from the command line, avoiding the need
of having to write R code to use it for the wrapped functionality. The sections available below

detail all the steps wrapped.

#### Garnett Exchange formats

- R CellDataSet (CDS) object (input/output)
- Tab-separated text file (output)

#### Garnett modules available

Table 1 below details all modules available as both a CLI component and as a Galaxy wrapper. The name of the module links to an active Galaxy instance where that can be used, and TS provides a second link to the module in the Galaxy Toolshed.
Each module is linked to one of the cli-layers to one or more of the relevant analysis areas: Clustering (C), Differential expression/Marker detection (DE-MD), Trajectories (T), Cell type alignment (CT), quality control (QC) and Dimensionality reduction (DR).

| Module | Description | cli-layer | Analysis areas |
|---|---|---|---|
| garnett_check_markers TS | Check marker file to filter out markers of suboptimal quality | garnett-cli | |
| garnett_classify_cells TS | Classify cells into cell types | garnett-cli | |
| garnett_get_feature_genes TS | Obtain a list of genes used as features in classification model | garnett-cli | |
| garnett_get_std_output TS | Get final output in standard format to allow for downstream analysis of predicted labels by tools of the EBI gene expression group's cell-types-analysis package | garnett-cli | |
| garnett_train_classifier TS | Train classifier based on marker gene list | garnett-cli | |

| | | |
|---|---|---|
| garnett_transform_markers[TS] | Transform marker files from Single Cell Expression Atlas format to that compatible with Garnett | garnett-cli |
| update_marker_file[TS] | Update marker file by filtering out suboptimal markers | garnett-cli |

### SCPred

From its website, SCPred:

> …is a general method to predict cell types based on variance structure decomposition. It selects the most cell type-informative principal components from a dataset and trains a prediction model for each cell type. The principal training axes are projected onto the test dataset to obtain the PCs scores for the test dataset and the trained model(s) is/are used to classify single cells.

SCPred is used as an R library, so R code needs to be written to use it directly. We have written
a CLI component named scpred-cli which allows its usage from the command line, avoiding the need
of having to write R code to use it for the wrapped functionality. The sections available below detail all the steps wrapped.

#### SCPred Exchange formats

- SingleCellExperiment (input/output)
- Tab-separated text file (output)

#### SCPred modules available

Table 1 below details all modules available as both a CLI component and as a Galaxy wrapper. The name of the module links to an active Galaxy instance where that can be used, and TS provides a second link to the module in the Galaxy Toolshed.
Each module is linked to one of the cli-layers to one or more of the relevant analysis areas: Clustering (C), Differential expression/Marker detection (DE-MD), Trajectories (T), Cell type

alignment (CT), quality control (QC) and Dimensionality reduction (DR).

| Module | Description | cli-layer | Analysis areas |
|---|---|---|---|
| scpred_eigen_decompose[TS] | Perform matrix eigen-decomposition; initialize object of scPred class | scpred-cli | |
| scpred_get_feature_space[TS] | Get feature space for training matrix | scpred-cli | |
| scpred_get_std_output[TS] | This method allows to export predicted labels in a standardised format, simplifying downstream analyses. | scpred-cli | |
| scpred_predict_labels[TS] | Make cell type predictions using trained model. | scpred-cli | |
| scpred_train_model[TS] | Train classification model | scpred-cli | |
| scpred_traint_test_split[TS] | CPM normalise and partition into train/test data | scpred-cli | |

### Cell-type-analysis

A suite of scripts for analysis of scRNA-seq cell type classification tool outputs. These scripts can be used both for evaluating the existing methods by running pipelines on labelled data and for analysing predicted labels for novel data sets.

#### cell-types-analysis Exchange formats

- Tab-separated text format (input/output)
- R `RDS` object (input/output)

#### cell-types-analysis modules available

Table 1 below details all modules available as both a CLI component and as a Galaxy wrapper. The name of the module links to an active Galaxy instance where that can be used, and TS provides a second link to the module in the Galaxy Toolshed.
Each module is linked to one of the cli-layers to one or more of the relevant analysis areas: Clustering (C), Differential expression/Marker detection (DE-MD), Trajectories (T), Cell type alignment (CT), quality control (QC) and Dimensionality reduction (DR).

| Module | Description | cli-layer | Analysis areas |
|---|---|---|---|
| ct_build_cell_ontology_dict[TS] | Create a mapping from labels to CL terms | cell-types-analysis | |
| ct_combine_tool_outputs[TS] | Combine predictions for single tool from multiple datasets | cell-types-analysis | |
| ct_get_consensus_outputs[TS] | Get consensus outputs across multiple tools | cell-types-analysis | |
| ct_get_empirical_dist[TS] | Get empirical distribution for tool performance table | cell-types-analysis | |
| ct_get_tool_perf_table[TS] | Get performance table for a list of outputs generated by various tools | cell-types-analysis | |
| ct_get_tool_pvals[TS] | Get p-values for tool performance metrics | cell-types-analysis | |
