## Supplemental Note 3 for "User-friendly, scalable tools and workflows for single-cell analysis"

### Galaxy tools

This part of the setup explains how to deploy all of our latest tools and workflows on a living Galaxy instance, for which you will need to know its URL, port and have an API key from an admin user that can install tools in the instance. This could have been achieved in different ways, but once the Galaxy instance is up, it becomes irrelevant how it was provisioned for the sake of adding our tools and workflows.

These instructions also adds third party tools that we have considered useful through the time that we have used this setup for production, exploratory analysis and trainings.

The most up-to-date version of this supplementary material will be available here.

#### Tools installation

Tools to be provisioned are mostly available through the main Galaxy Toolshed within our ebi-gxa user. There are additional third party tools that we provision as part of this process. All versions of each tool are installed, which means that workflows created at different points in time will work with the tools that they were originally created with. Normally, they user should be able to upgrade the tools in a workflow through the interface without much issues, but this design guarantees reproducibility.

Requirements:

- Conda
- Conda environment with Galaxy ephemeris. This can be achieved via:

```
conda create -n galaxy-ephemeris -c bioconda ephemeris
```

- A Galaxy API key for an administrator user in the instance where you would like to install the tools.
- Inside `tool_conf.xml` Galaxy config file, place the following content (this is not necessary if you are using the helm/k8s setup):

```
<label id="single_cell" text="Single Cell RNA-Seq Tools"/>
<section id="hca_sc_get-scrna" name="Get scRNAseq data">
 </section>
<section id="hca_sc_seurat_tools" name="Seurat">
 </section>
```

```
<section id="hca_sc_sc3_tools" name="SC3">
 </section>
<section id="hca_sc_scanpy_tools" name="Scanpy">
 </section>
<section id="hca_sc_monocle3_tools" name="Monocl3">
 </section>
<section id="hca_sc_scmap_tools" name="SCMap">
 </section>
<section id="hca_sc_sccaf_tools" name="SCCAF">
 </section>
<section id="hca_sc_utils_viz" name="Single Cell Utils and Viz">
 </section>
<label id="rna_seq_label" text="Bulk RNA-Seq Tools"/>
<section id="rna_seq" name="RNA-Seq">
 </section>
<section id="gxa_util" name="Util">
 </section>
```

You can re-arrange the order of sections or their names, but their identifiers should be kept.

Please note that tools will only appear in the Galaxy instance if a section identifier matches between the above and
the content of the .lock files in https://github.com/ebi-gene-expression-group/galaxy-gxa-tools-setup.

To be executed in bash

```
# clone our tool definition file repository
git clone https://github.com/ebi-gene-expression-group/galaxy-gxa-tools-setup
# activate the ephemeris conda env, provided it was created as shown in the require
conda activate galaxy-ephemeris
cd galaxy-gxa-tools-setup
./scripts/install_tools.sh http://you-galaxy-url:port/ <GALAXY_ADMIN_API_KEY>
```

The above github repository is routinely updated with newer revisions of the tools, so if you want to keep
up-to-date, you might want to run this routinely.

### Conda package installation

Any of the conda packages mentioned in the Tools Supplementary note can be installed, once conda is available,
through:

```
conda create -n <name-of-an-environment> <name-of-conda-package>
```

For example, for scanpy-scripts:

```
conda create -n my-env-with-scanpy scapy-scripts
```

The bioconda entry for that example is available here.

### Access through containers

Any of the conda packages mentioned in the Tools Supplementary note can be pulled/run, once docker is available, through (specifying the tag):

```
docker pull quay.io/biocontainers/scanpy-scripts:<tag>
```

you can find available tags for that container here.
