## Supplemental Note 4 for "User-friendly, scalable tools and workflows for single-cell analysis"

### Running Hinxton Singe Cell Galaxy Setup

The most up-to-date version of this document is available here.

Running the galaxy setup with our tools and workflows can be done in various ways, from easier to more complex (but more powerful):

- Directly through https://humancellatlas.usegalaxy.eu/. This is a Galaxy instance that is ready and hosted by the Frieburg Galaxy group on a DEN.BI funded HPC system with access to >7,000 cores and ~50 TB of RAM through ~200 nodes, as of the time of this writing.
- On an existing Galaxy instance where you have administrative rights or where you can ask administrators to install tools that you need. Ask the administrator to follow these instructions (or the direct tools installation supplementary material) to install our tools and workflows there.
- On your own machine through Kubernetes (k8s). This entails having k8s installed on your laptop/desktop machine. This will allow you to run most tools, but of course performance is degraded compared to deployment in an HPC or cloud solution due to having less hardware available (less CPUs, RAM, disk, etc).
- Deploying on a cloud provider through k8s. This will vary from cloud provider to cloud provider, but the essential steps are:
    - Deploy a k8s cluster
    - Deploy a shared file system that is accessible from that k8s cluster or make sure that the k8s cluster has a storage class for ReadWriteMany storage. This will give rise to your Galaxy Persistent Volume Claim (PVC) or the storage class to use for Galaxy.
    - (Optionally) Deploy a file system that can be accessible at least from one node for Postgresql, or make sure that you have access to a storage class for this. This will give rise to a Postgresql PVC or storage class.
    - Obtain authentication (usually a `kube-config.yaml` file) for the k8s cluster so that you can use `kubectl`.
    - Deploy the Galaxy helm chart against that cluster.
    - Download this config file and modify it following the instructions in all the TODO comments inside (so, Galaxy PVC, admin email and Postgresql PVC). You can find more variables to adjust here.
    - Make sure that you have helm and kubectl binaries installed.
    - Obtain the helm chart by running:
      `helm repo add galaxy-gvl https://raw.githubusercontent.com/cloudve/helm-charts/master`
    - Install the chart with your modified config:
      `helm install -f hsciap-20.01-helm-galaxy-v3-modified.yaml galaxy-gvl/galaxy`

- Galaxy will be available at any of the k8s nodes public IPs, at port 30700. Go there, register with one of the set administrator emails and in the user preferences, generate an API key, to be used in the next step for loading tools.
- Deploy the tools as per instructions.
  As an example of the above, these instructions allow to deploy the setup in AWS.
