## Supplementary Table S1 for "User-friendly, scalable tools and workflows for single-cell analysis"

Table S1: Command line argument layer for each tool, the analysis areas covered and with links to the different packaging generated: Conda package (C) and Docker Containers (DC).

| Tool | cli-layer | Packaging | Analysis areas |
|---|---|---|---|
| Scanpy | Scanpy scripts | C DC | clustering, dimensionality reduction, differential expression. |
| Seurat | Seurat scripts | C DC | clustering, dimensionality reduction, differential expression. |
| SC3 | SC3 scripts | C DC | clustering, dimensionality reduction, differential expression. |
| Scater | Scater scripts | C DC | clustering, dimensionality reduction, differential expression. |
| scMap | scMap cli | C DC | cell type alignment |
| scPred | scPred cli | C DC | cell type prediction |
| Garnett | Garnett cli | C DC | cell type prediction |
| Monocle3 | monocle scripts | C DC | clustering, trajectories. |
| UCSC CellBrowser | None needed | C DC | interactive visualisation |
| SCCAF | Contributed package includes cli | C DC | clustering |
| SCEasy | Contributed packaged includes cli | C DC | Format conversion |
| DCP Matrix Service Client | Contributed package includes cli | C DC | selection & aggregation |
